# License to cut: Smart RNA guides for conditional control of CRISPR-Cas9

**DOI:** 10.1101/2022.10.26.513620

**Authors:** Alexandre Baccouche, Arman Adel, Nozomu Yachie, Teruo Fujii, Anthony J. Genot

## Abstract

The Cas9 enzyme is a programmable endonuclease, whose target sequence is directed by a companion RNA guide. Cas9 and RNA guides have revolutionized biology, enabling facile editing of the genome in almost all organisms. Controlling where and when Cas9 and the guide operate is indispensable for many fields ranging from developmental biology to therapeutics, but it remains a challenge. Most methods focus on controlling Cas9 with physico-chemical means (which lack finesse, precision or multiplexing), or transcriptional tools (which are slow and difficult to design). Rather than directly engineering Cas9, engineering the RNA guide itself has emerged as a more general and potent way to manage the activity of Cas9. Here we report smart RNA guides that are conditionally activated by the presence of a specific RNA opener. Contrary to most previous approaches, the design affords ample freedom as spacer and the opener are independent. We demonstrate this flexibility by operating SmartGuides activated by a panel of miRNA relevant for human health, and by composing SmartGuides in Boolean logic circuits. Lastly, we test the SmartGuides in mammalian cells - validating the basics tenets of the design, but also highlighting the challenges that remain to be lifted for in-vivo operation.

## Introduction

Cas9 is a programmable endonuclease that is directed by a RNA guide to cut a target DNA sequence^1–3^. The ease of programming Cas9 has enabled a range of new applications, ranging from base and gene editing to genetic and epigenetic regulation^4,5^. As Cas9 has expanded to all walks of life sciences, so has the need for finer control of its activity^6^. Once Cas9 and a RNA guide are present in a cell, the tandem will inevitably cleave its DNA target, irrespective of the type, environment, internal state or developmental stage of the cell. But mistimed, misplaced or mismatched cleavage is detrimental, and researchers are looking for methods to decide when, where and how Cas9 is activated.

Temporal control of Cas9 is desirable for instance in developmental biology because premature knockout of a vital gene causes embryonic lethality^7–9^. Controlling the timing of Cas9 is also important for improving the efficacy and safety of gene editing. Prolonged activity of Cas9 after germline editing has been linked to mosaicism in embryos^10,11^. Synchronizing Cas9 with specific cellular cycles promotes homology directed repair over non-homologous end joining - improving the precision of gene editing^12,13^.

Spatial control of Cas9 is also needed in multicellular organisms to confine its action to specific tissues or organs. The conventional method for conditional knockout in mice (gene floxing controlled by a cell-specific transcription factor^14^) is laborious and delicate, and only a handful of institutions can establish and maintain a repository of floxed mice. Gene knockout would be considerably simplified if the action of Cas9 could be conditioned to the presence of a transcript that is specific to a certain cellular type. More generally, Cas9 bears strong promises as a therapeutic tool, but concerns linger about off-target effects - limiting the tolerable dose of Cas9 that can be administered^15^. A tool that mutes Cas9 in healthy cells and activates it in diseased cells could help to administer stronger doses of Cas9 - improving its efficacy

Conditioning Cas9 to specific cellular states may enable new applications ranging from the scale of a single-cell to a whole population ^16–19^. At the single-cell level, cellular recorders log molecular events into DNA memories with Cas9. But their generality remains limited without a tool that couples a specific cellular event (e.g. the transcription of a given mRNA) to the editing of the memory register by Cas9. The majority of events that have been recorded so far with Cas9 are either random mutations by Cas9 (which is the basis for reconstructing cell lineages^20^), or externally triggered signals (small molecules, light…^21^). Conditioning Cas9 to the presence of specific endogenous RNA (messenger RNAs, long noncoding RNA, microRNA…) would widen the repertoire of recordable events, and enable the time-resolved recording of transcription profiles.

At a much larger scale, gene drives based on Cas9 could remodel exponentially fast the genetic makeup of a whole population^22^. In the future, gene drives could, for instance, eradicate mosquito-borne pathologies, but for the moment they raise serious environmental issues, because a copy of Cas9 is left lurking in the genome of the mutated population, opening the unpalatable prospect of genomico-ecological catastrophes. These concerns would be alleviated if this genetic circuit could carry a kill switch, an instruction that would force Cas9 to erase itself from the genome after a given number of generations has passed^23,24^. This self-erasure of Cas9 could be achieved by combining a heritable DNA ledger (that accumulates mutations at a constant rate), and a Cas9 that is conditioned to knock out its own gene once the ledger has been fully mutated.

Engineers have followed four complementary routes to control Cas9: chemical engineering, protein engineering, genetic engineering and RNA engineering. Chemical engineering designs or discovers chemical compounds that inhibit Cas9, such as Anti-CRISPR proteins^25,26^. This goes hand in hand with protein engineering, which designs variants of Cas9 that are susceptible to chemical signals or physical signals (e.g. light or temperature)^27,28^. But physico-chemical control of Cas9 lacks single-cell precision and multiplexing. The signal can be toxic to cells, and its delivery is a limiting factor: optical methods are hindered by the low penetrability of visible light into deep tissues, whereas chemical methods are limited by the challenge of timely and localized delivery. Most importantly, those methods are unconditional. The signal is exogenous and activates Cas9 irrespective of the state of the host cell. Many applications such as cellular recording call for an endogenous control of Cas9, where the signal inducing Cas9 is produced by the cell itself rather than by an external operator.

Genetic engineering - one of the main tools of synthetic biology - offers this endogenous control and has been explored for the CRISPR-Cas9 system. It places the transcription or translation of the Cas9 gene, an RNA guide, or even an anti-CRISPR gene, under the control of a factor that is only expressed under certain conditions^29^, or inside specific cells ^30–32^. But transcriptional control of Cas9 suffers from some of the classical drawbacks of genetic circuits: delayed response, poor predictability, paucity of actionable transcription factors and overloading of the host’s machinery^33^.

Rather than engineering the transcription, translation or activity of Cas9, a promising alternative is to engineer the RNA guide itself. Firstly, RNA strands are far simpler to sequence, express or modify than proteins. Secondly, RNA sensors can be installed in RNA guides to interface them with the transcriptome - providing access to a rich trove of cellular information. A natural approach to control the RNA guide is to sequester its spacer domain with a complementary RNA sequence, which is removed in the presence of a specific inducer - activating the guide^34–36^. Another approach is to stabilize or destabilize portions of the RNA guide with ligands^37^, or to process the guide with an enzymatic machinery. Wang et al. designed a precursor of an RNA guide which is constitutionally inactive. But in the presence of a specific microRNA, it is processed by the RISC machinery into a competent RNA guide^38^. This opens numerous possibilities as miRNA are overexpressed in many tissues and pathologies, which make them ideal biomarkers to infer the state or type of a cell. However this method is not universal: it is limited to organisms like plants or animals with a miRNA machinery (which prokaryotes lack), and is only applicable to miRNA (excluding the vast majority of the transcriptome). Lastly, it is unclear if the method can be scaled up to meet the multiplexing needs of methods like transcriptional recording. In the current design, the precursor guide (∼100 bp) needs a large promoter (∼2000 bp). Assembling vectors for dozens or hundreds of miRNA would be a herculean effort, and would undoubtedly burden the host’s RNAi machinery.

In parallel, several groups have leveraged the tenets of DNA nanotechnology to allosterically regulate RNA guides with a general and non-enzymatic mechanism: toehold-mediated strand displacement^39,40^. The mechanism is based solely on Watson-Crick base pairing and does not require external factors (such as a Dicer protein), so it could be deployed in any organism. Most designs start by sequestering the spacer of the RNA guide with a complementary RNA domain, which abolishes the activity of Cas9. The presence of a trigger RNA initiates a toehold-mediated strand displacement, and releases the spacer. Some of the initial designs imposed a strong homology between the spacer and the trigger RNA. This excludes the most sought-after applications where the trigger RNA and the DNA target are endogenous and unrelated. For instance, for targeted gene editing,the trigger could be a miRNA marker of a given cellular type, while the DNA target is the loci of the gene to be edited. Subsequent designs have improved orthogonality between the trigger RNA and the spacer^41–43^. Li and colleagues proposed an elegant design in which a tail appended to the guide folds back and binds to the loop of the upper stem, preventing the guide from interacting with Cas9^44^. Addition of a target RNA removes this folding and restores functionality. Interestingly, this design removes homology constraints and in principle could link unrelated input RNA and output DNA. However the length of the input must be in the 50 nt range, which prevents operation with short RNA like miRNAs.

Here we report smart RNA guides for Cas9 that are allosterically modulated by a small and arbitrary RNA input. We modulate the affinity of the guide for Cas9 by controlling the folding of a crucial secondary structure: the nexus. Using an *in vitro* transcriptional circuit, we demonstrate orthogonal and multiplexed regulation of the SmartGuides with a panel of miRNAs relevant for human health. We then demonstrate higher-order computation with the SmartGuide, such as non-monotonic Boolean logic. Lastly, we coupled the SmartGuide to a base-editor and explored their activity in mammalian cells - laying out challenges that will need to be overcome for a future deployment in complex organisms.

## Results

We benchmarked the SmartGuides *in vitro* with a transcriptional circuit based on a fluorescent reporter for Cas9^45^. Compared to the conventional workflow for assessing the activity of RNA guides *in vitro* (overnight transcription, gel purification, cleavage assay and finally gel electrophoresis^1^), this one-pot assay not only reduces the hands-on time from a few hours to a dozen minutes, but also provides quantitative, multiplexed and time-resolved kinetic measurements - allowing us to test ∼300 designs of guides over the course of two years. A one pot assay is also a more realistic model of the cell than a conventional cleavage assay, because all processes occur concurrently (transcription and cotranscriptional folding of the guide, opening of the guide by the miRNA, loading of the opened guide by Cas9, and finally cleaving).

The key element of our control strategy is the nexus of the RNA guide, a short hairpin located near the first stem of the scaffold^46^. In the Cas9:guide complex, the nexus makes extensive contacts with Cas9, and sits at the interface of α-helical lobe (used for recognition) and the nuclease lobe of Cas9 (used for cleavage)^47,48^. This makes the nexus an ideal target for allosteric regulation, because small changes to the nexus have an outsized influence on the activity of Cas9. For instance, mutating the two nucleotides that hold the nexus suppressed the hairpin structure and completely abolished the activity of the guide^46^.

Informed by these structural and functional insights about the criticality of the nexus, we installed two modifications on the canonical RNA guides to control the folding of the nexus. Firstly, we appended a tail fully complementary to the cognate miRNA. Secondly, we mutated the sequence of the nexus and the bridge - while preserving the secondary structure of the nexus - to make this region partially complementary to the tail and form a hairpin. It prevents the folding of the nexus and the loading by Cas9, resulting in a guide that natively folds in an inactive state. The presence of a cognate miRNA rescues the activity of the SmartGuide. The miRNA binds to the sequestering hairpin by a long toehold, unzips it by toehold-mediated strand displacement (ref), and releases the nexus. This guide presents a canonical - though mutated - secondary structure which is loaded by Cas9 for enzymatic processing.

We tested two designs of the guide: one with a toehold located outside the blocking hairpin (external toehold), and one with a toehold located inside (internal toehold). Both SmartGuides display. In the ON state (opener present), the fluorescence of the reporter rises in about 60-120 min to ∼50-100% of the fluorescence of the control guide. Various factors could explain this discrepancy between control guides and SmartGuides : secondary structures in the miRNA or the SmartGuide, lower transcription efficiency of the SmartGuide, or altered contacts of the mutated nexus of the SmartGuide with Cas9. In the OFF state of the guide (no opener present), the reporter displays a leak activity corresponding to ∼ 10-20% of the ON state. Interestingly, the time trace informs about the possible origin of the leak. If the leak was caused by a subpopulation of guides that were incorrectly transcribed or folded (thus failing to turn OFF), we would observe a slow upward drift of fluorescence which would eventually reach the same level as the control (because the guides are continuously transcribed over the course of the experiment, and their concentrations will eventually vastly exceed that of the reporter). Rather we observe a quick jump in fluorescence followed by a plateau. This suggests that the leak may instead be caused by a faulty subpopulation of reporters, which is incorrectly synthesized or assembled.

We then benchmarked the SmartGuides against a panel of 6 representative human miRNAs. They are are enriched in endothelial cells (mir-21^49^), hypertrophic scarring tissues (mir-98^50^), cancer cells (mir-451^51^), psoriasis skin cells (mir-31^52^), neutrophils following traumatic injury (mir-3945^53^) or in the liver (mir-122^54^). When the SmartGuides and the miRNA matched (traces on the diagonal), the fluorescence of the reporter showed strong and fast activation. When they were mismatched, activation remained negligible (at ∼5-10% of the ON level). Half of the ON guides in the panel displayed a kinetic and end-point fluorescence similar to the control (namely mir-98, mir-451 and mir-21). For the other half, the kinetic was slower than the control, and finished at a lower level (mir-3945, mir-31 and mir-122). However, all guides showed robust levels of activation on a timescale compatible with in-vivo application (minutes to hours).

In view of making Cas9 safer and more precise, we composed SmartGuides to demonstrate Boolean computations. In real-world settings, simply testing the presence or absence of a single miRNA may not be sufficient, and one may condition Cas9 on the presence of a panel of miRNAs that form a signature of a cellular state. This would improve the specificity of Cas9, by ensuring that it is only activated inside cells or in response to situations that need it.

We first installed a NOT gate in the guide by mutating the hairpin loop after the upper stem. The mutated loop binds to the target RNA, which disrupts the secondary structure of the guide and abolishes its function (Fig. 3b). We then combined conditional activation and inhibition, and computed a NOT AND gate. The gate only fires in the presence of target input and in the absence of another one (Fig. 3b). Lastly, we computed a XOR gate, one of the most complex Boolean functions on two inputs. The circuit comprises two symmetrical guides, each computing a NOT AND gate on the same pair of inputs. As expected, the circuit only fires when one - and only one - input is present. The activation level, while robust, is noticeably lower than the control guide. Lastly, we multiplexed two SmartGuides with distinct spacers, demonstrating their concurrent and individualized operation in the same mixture (Fig. 3b). This multiplexing is an added benefit compared to physical or chemical methods to control Cas9 (e.g. with small molecules, light or temperature), as it is difficult to expand the repertoire of their actionable signals.

Encouraged by the robustness and versatility of the SmartGuides *in vitro*, we tested them in human cells. We placed a Cas9 base editor^4^ under the control of a SmartGuide - using a miRNA or a synthetic RNA as input (Figure 4). When the guide is ON, it directs the base editor to edit a base in a mutated GFP gene, rescuing its activity and causing the cell to fluoresce green. All components of this genetic circuit (the input RNA, the guide, the GFP gene and the Cas9-Target AID fusion gene) are genetically encoded in plasmids and transfected in the cells. Surprisingly, we found that when operated *in vivo*, Cas9 did not tolerate mutations in the guide that it tolerated *in vitro*. A control guide, lacking the blocking tail but harboring in the nexus and the linker, failed to light cells up (Fig. 4c left). We surmised that some mutations in the nexus disrupted contacts with Cas9, thus lowering the affinity of the guide with Cas9. While this lower affinity may not have been problematic *in vitro*, it has been hypothesized that *in vivo*, the RNA guide must compete with non-specific RNA for binding with Cas9, and thus require a strong affinity to overcome this competition^45^. To test this hypothesis, we rescued mutations in the nexus, but not the linker of the SmartGuide. In turn, this required us to modify the design of the SmartGuide, because the unmutated nexus breaks the complementarity of the tail with the input RNA. We moved the toehold internally, putting it inside the loop and at the base of the hairpin neck. Opening of hairpins with internal toeholds has been studied *in vitro*55, and though it progresses at a slower pace than for external toeholds, it remains sufficiently fast for biological timescales. We tested the design with an internal toehold *in vitro* and confirmed that it worked satisfactorily (Fig. 1c).

**Figure 1:**
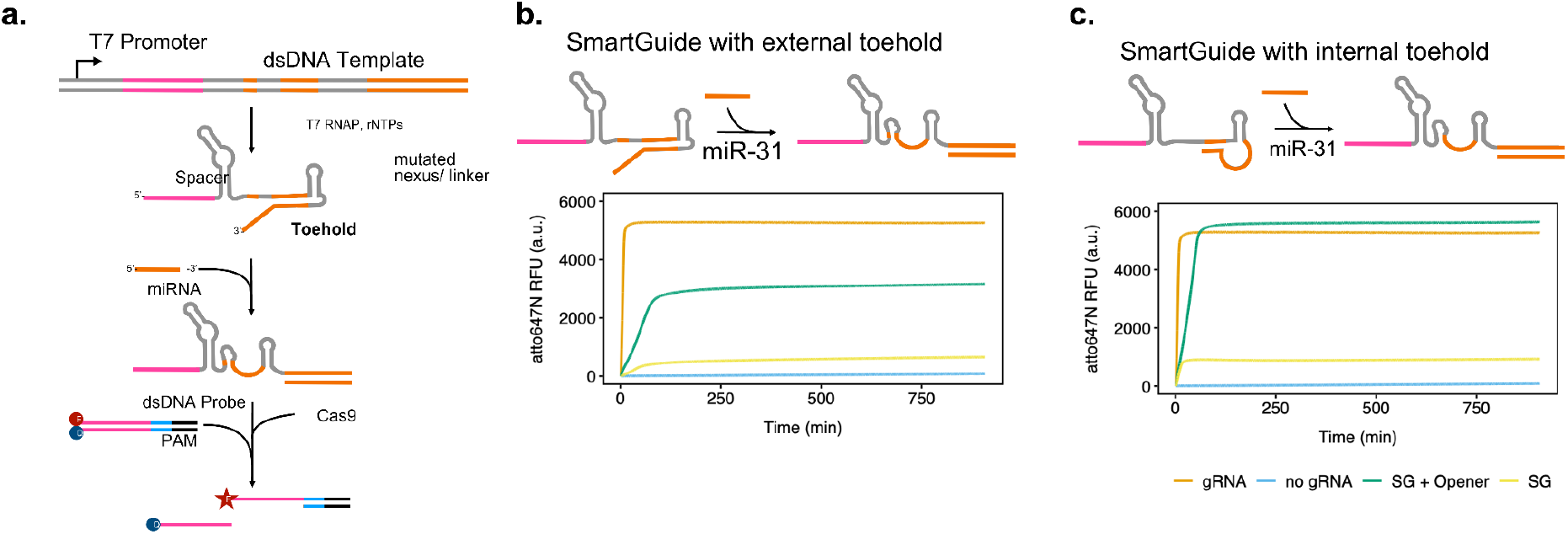
Smart RNA guides for conditional control of Cas9. a. The SmartGuides (SG) are benchmarked with a transcriptional circuit. The guide is transcribed from a dsDNA template and bears a tail in 3’ that folds back onto the nexus and linker domain - preventing loading of the guide by Cas9. But the presence of a target miRNA opener rescues the nexus by invading the terminal hairpin through a single-stranded toehold domain. This invasion releases the nexus and exposes a canonical guide that is loaded by Cas9. This guide:Cas9 complex invades and cleaves a fluorescent reporter, generating an increase in fluorescence. The process is one-pot and all reactions occur concurrently. b. Testing of a SmartGuide with a toehold located outside the terminal hairpin loop (external toehold). The plot shows the fluorescence level of the beacon. gRNA is the control trace (canonical guide that is constitutively ON). c Testing of a SmartGuide with a toehold located inside the terminal hairpin loop (internal toehold). gRNA is the control trace (canonical guide that is constitutively ON).

**Figure 2:**
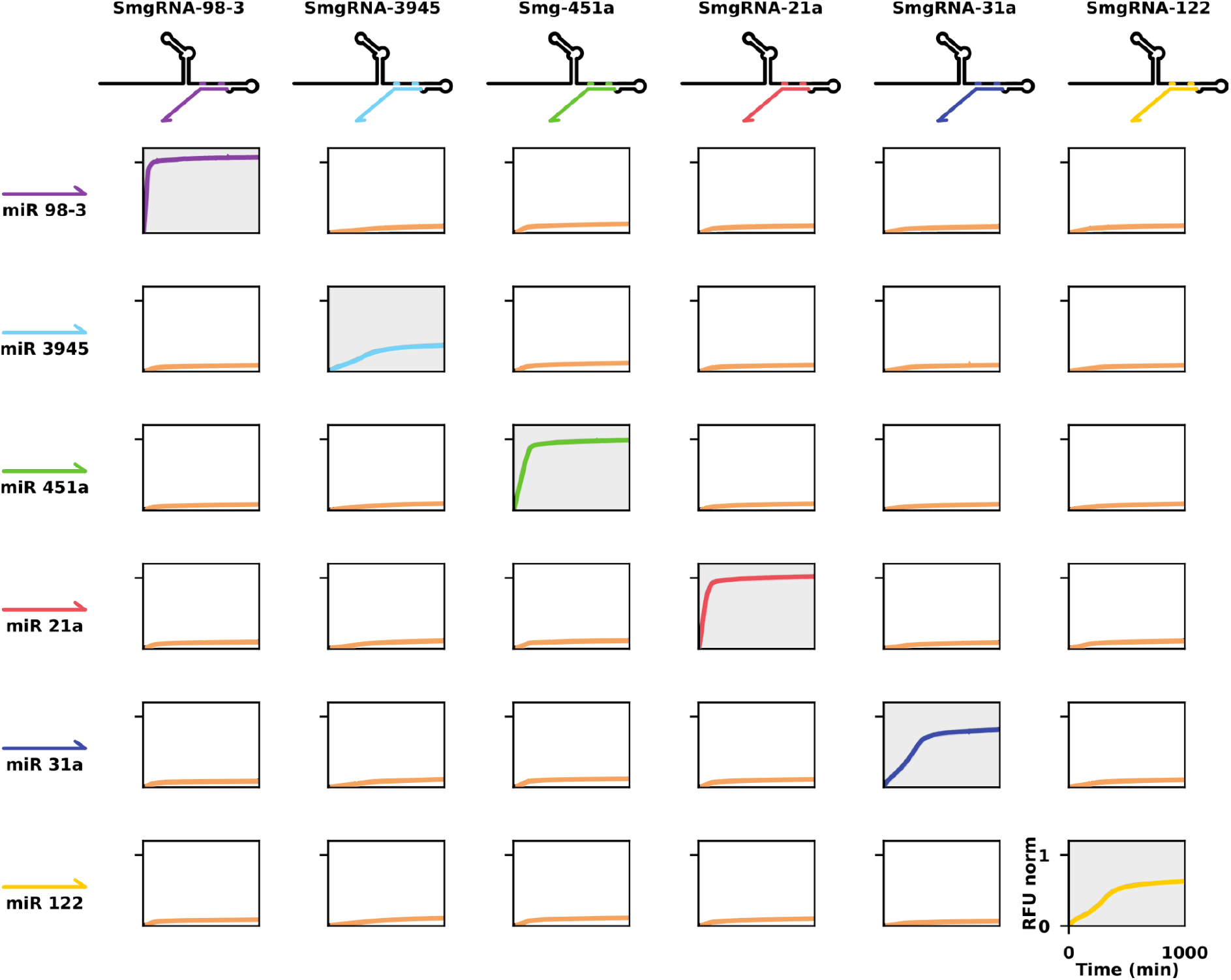
Activation of SmartGuides against a panel of miRNA, measured in a one-pot assay. Each plot shows the fluorescence timetrace of the reporter following mixing of one guide (top row) against one miRNA input (left column). Traces on the diagonal correspond to ON-target activation, while traces off the diagonal correspond to OFF-target activation. The traces are normalized with respect to their control guide (i.e. the SmartGuide lacking the blocking tail but harboring a mutation in the nexus).

**Figure 3:**
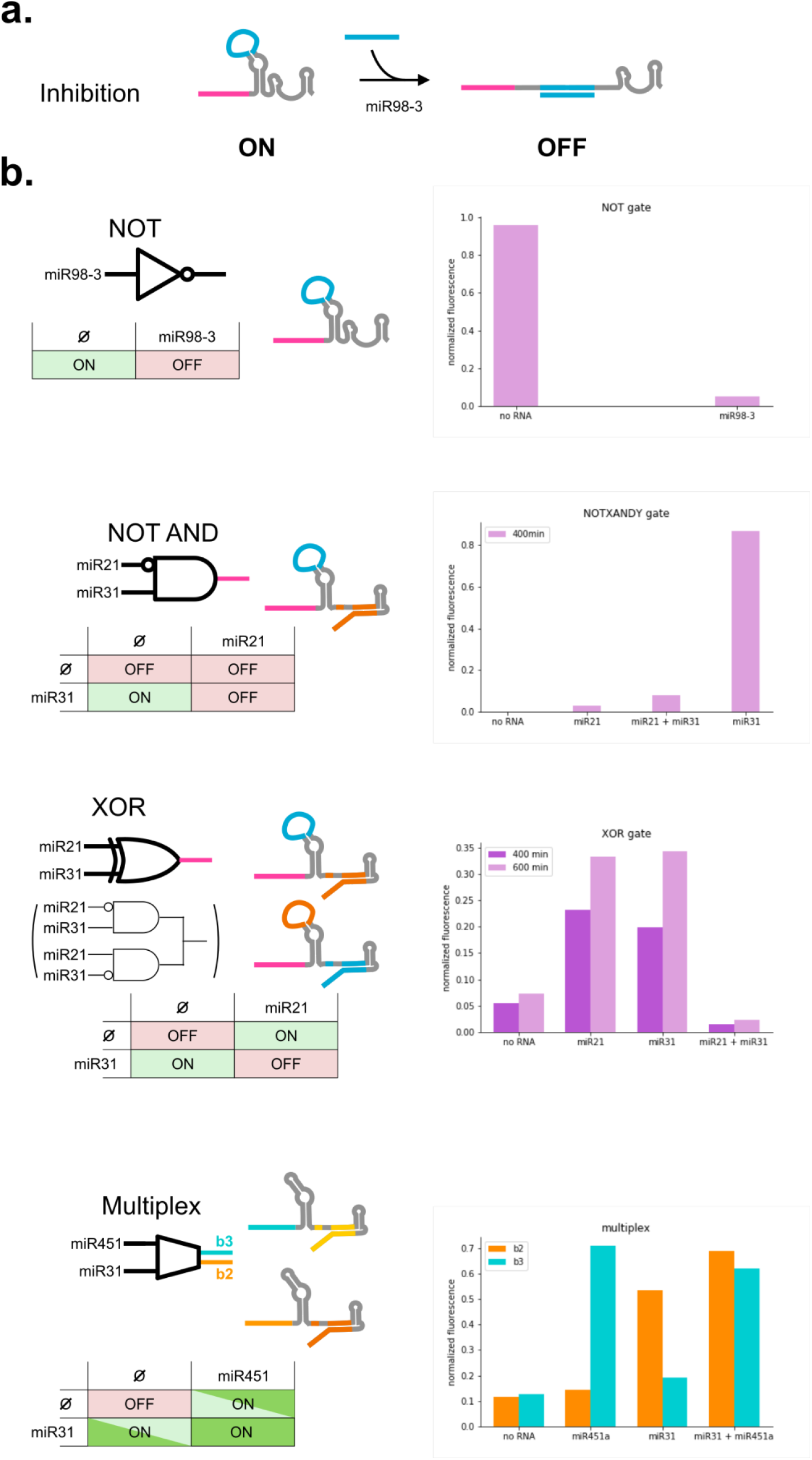
Logical operation and multiplexing of the SmartGuides. **A** Scheme for computing a NOT gate by mutating a RNA guide. The guide is constitutively ON. Addition of the input RNA opens the stem loop (which has been appropriately mutated to bind the RNA) and abolishes functionality of the guide. **B** Computation of various logic gates by composing the conditional activation of the SmartGuide with the NOT gates. **C** Multiplexing: orthogonal operation of two SmartGuides, each with its own opener, spacer and fluorescence reporters. (Note that we adjusted the readout time according to the composition of the logic circuit.). The fluorescence level is defined with respect to the maximum level of fluorescence reached by the control.

**Figure 4:**
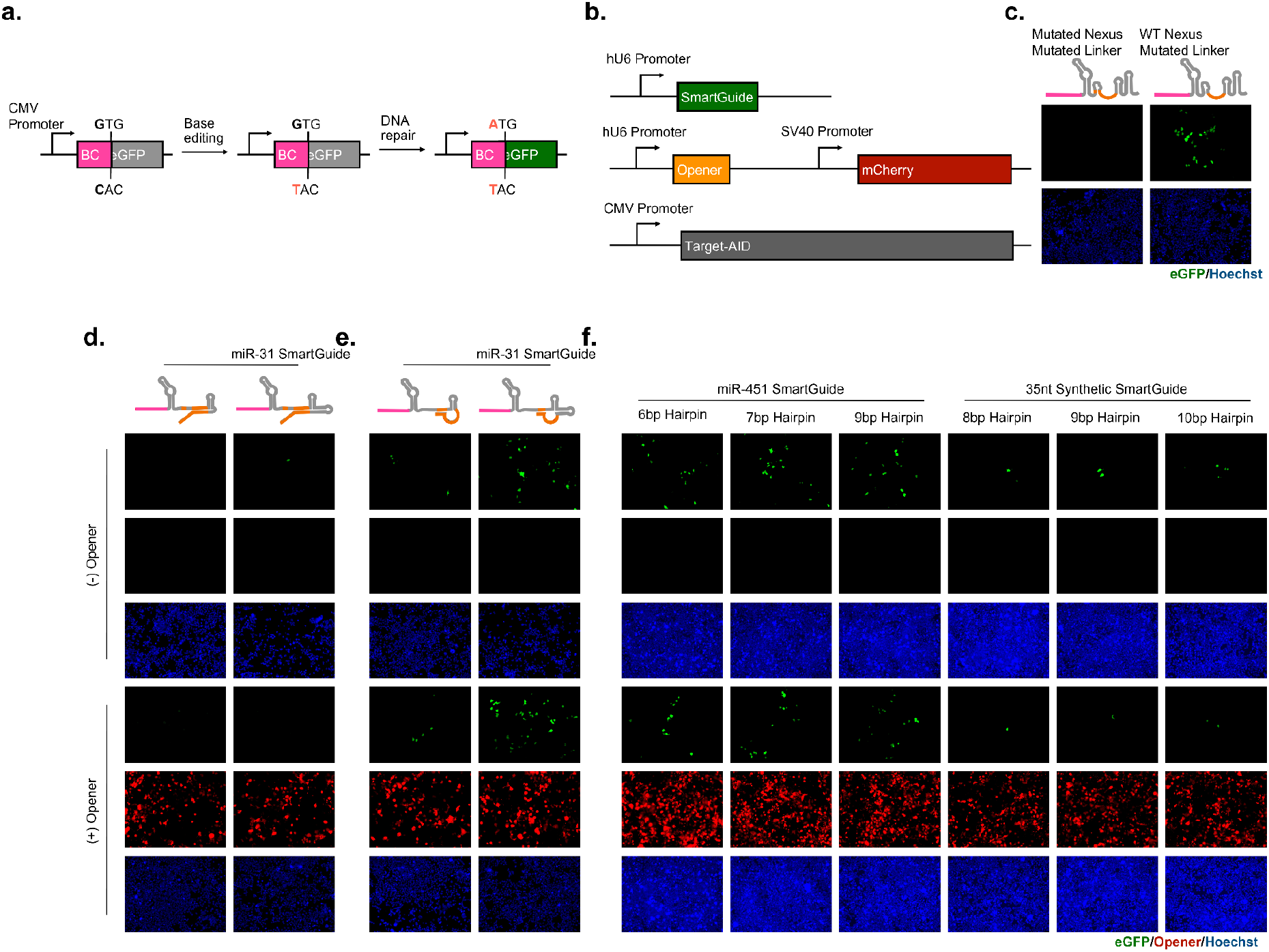
Arman’s In-vivo section. a) CRISPR-Cas9 base-editing reporter system b) Gene architecture of SmartGuide, small RNA opener and Target-AID cassettes c) gRNA activity for the original miR-31 responsive SmartGuide and redesign without nexus mutations in the absence of the toehold sequence d) Fluoroscope images of HEK293Ta base-editing reporter cells after transfection with miR-31 sequence responsive SmartGuide with Target-AID in the presence or absence of a short opener sequence e) Fluoroscope images of HEK293Ta base-editing reporter cells after transfection with redesigned miR-31 sequence responsive SmartGuide with Target-AID in the presence or absence of a short opener sequence f) Fluoroscope images of HEK293Ta base-editing reporter cells after transfection with redesigned miR-451 or synthetic sequence responsive SmartGuide of varying hairpin length (in orange) with Target-AID in the presence or absence of a short opener sequence.

We then attempted to operate the SmartGuide by triggering them in presence of microRNA. Satisfyingly, we observe that the control ON guide has a basal activity similar to that of the wild type guide (Fig. 4c right) - validating our hypothesis about the *in vivo* tolerance to mutations in the linker but not in the nexus. Additionally, the SmartGuide in its OFF state displays little fluorescence (Fig. 4e left), validating the tight control exerted by the tail on the nexus. However, expression of the opener RNA led to a modest increase of fluorescence - indicating that the SmartGuides do not fully switch ON in presence of their target. We tested various lengths of hairpins with two different openers, but could not observe a significant increase of green fluorescence in presence of the opener.

This lack of switching *in vivo* is not likely due to design: guides designed to be in the ON states are ON, and guides designed to be in the OFF state are OFF. We surmise that the failure to completely switch *in vivo* could be due to the complexity of nuclear biology: the guide and the opener may be expressed far away in the nucleus and never meet, or they could be exported outside of the nucleus and meet in the cytoplasm, but never traffic back to the nucleus. In addition, the secondary structures of small RNA are known to be highly regulated ^56,57^. Further studies with live cell imaging could illuminate the fate of the guide and the opener RNA. Studies in prokaryotes are also warranted to ascertain the workings of the SmartGuide in a living organism.

## DISCUSSION

We have reported smart RNA guides for Cas9 that are conditionally activated by the presence of an arbitrary RNA. The spacer and the input RNA are unrelated, which expands the range of circuits that can be designed. While we have used miRNA as a proof-of-principle, our design does not rely on miRNA-processing machinery, and could in principle apply to any RNA sequence. Operation of the SmartGuide in mammalian cells confirms the ON and OFF states of the SmartGuide, but the switching between those states remains incomplete and warrants further studies.

In the future, such SmartGuides may perform programmatic transactions on the genome based on the state of the in which they operate. This could make gene therapy safer and more potent, but also enable the recording of transcription events in a DNA memory.

## DATA AVAILABILITY

The data that support the findings of this study are available on request from the corresponding author.

## Methods

### Plasmid cloning

Plasmid cloning was carried out by conventional cloning methods such as PCR amplification, restriction enzyme digest, ligation and golden gate assembly. PCR amplification products were all generated using Phusion High-Fidelity Polymerase (NEB) and all restriction digests were carried out using NEB High-Fidelity restriction enzymes unless stated otherwise.

#### Lentiviral Base-editing reporter

Lentiviral C→T base-editing reporter plasmid (LV-CS-sg1-GTG-EGFP-Pur) reporter cassette was constructed by amplifying the eGFP gene fragment from pLV-SI-112 (Addgene 131127) using the primer set AA-sg1-BC-CS-eGFP-FW/AA-eGFP-BamHI-RV followed by restriction enzyme digest by EcoRI/BamHI of the eGFP fragment and pLV-SI-112, respectively. The digested fragment was then cloned into the EcoRI and BamHI sites of the pLV-SI-112 using T4 DNA ligase (Nippon Gene).

#### SmartGuide plasmid backbone and Golden Gate cloning of SmartGuides

The PB4 plasmid backbone used to construct the SmartGuide plasmid backbone was a generous gift from Dr. Yangming Wang of Peking University. The SmartGuide backbone plasmid (PB4-sg1-scaf-cloning) was constructed by first generating the insert fragment by fusion PCR: The hU6 promoter fragment was generated by amplifying the U6 promoter with primer set AA-MluI-U6-gib-FW/AA-U6-sg1-scaf-short-RV from the plasmid backbone pSI-359 (Addgene 131131) and the dsRed filler fragment was generated by amplifying a portion of the dsRed gene from CMVp-dsRed2-Triplex-HHRibo-gRNA1-HDVRibo-pA (Addgene 55201) with primer set AA-sg1-scaf-short-filler-FW/AA-filler-SpeI-gib-RV in separate PCR reactions. After gel purification of each fragment, the second PCR reaction was carried out with equimolar amounts of the hU6p and dsRed fragments to a total input mass of 10ng and using primer set AA-MluI-U6-gib-FW/AA-filler-SpeI-gib-RV. The fusion PCR product and PB4 backbone were then respectively digested by KpnI/SpeI and the fusion PCR product was cloned into the PB4 backbone using T4 DNA ligase (Nippon Gene).

Control scaffold sequences and SmartGuide toehold sequences were cloned into SmartGuide backbone cassette by oligonucleotide annealing and Golden Gate Assembly (Supplementary Methods).

#### Small RNA Opener plasmids

In order to construct the SmartGuide opener plasmid, first an mCherry gene was amplified from a pcDNA3.1 plasmid (pSI-177) using the primer set AvrII-1x-AUG-XFP-FW/BstB1-mCherry-RV. The mCherry gene fragment and the backbone pcDNA3.1 plasmid (pSI-166) were then respectively digested by AvrII/BsrGI and then the mCherry fragment was cloned into the pcDNA3.1 backbone using T4 DNA ligase (Nippon Gene). Following this, the opener sequences were cloned in by amplification of the hU6p promoter from pSI-359 and then the opener fragment and backbone plasmid were respectively digested by MluI/XbaI and the opener fragment was cloned into the backbone plasmid using T4 DNA ligase (Nippon Gene).

### Cell culture

HEK293Ta cells were purchased from GeneCopoeia and maintained in Dulbecco’s Modified Eagle’s Medium (DMEM) (Sigma) supplemented with 10% fetal bovine serum (FBS) (Thermo Fisher Scientific) and 1% penicillin–streptomycin (Sigma) at 37 °C with 5% CO_2_. Cells were routinely tested for mycoplasma contamination by nested PCR using culture medium as a template.

### Generating base-editing reporter HEK293Ta cell line

The C→T base-editing reporter HEK293Ta cell line was generated by lentiviral transduction. HEK293Ta cells were first seeded in one well of a 6 well plate at a seeding density of 3×10^5^ cells in 2ml of DMEM 1 day prior to transfection. Plated cells were then transfected with 489 ng of lentiviral plasmid, 366ng of psPAX2 (Addgene 12260), 122ng of pMD2.G (Addgene 12259) and 9.38 μl of 1 mg/ml polyethyleneimine MAX (PEI) (Polysciences) in 200 μl Opti-MEM (Gibco). 18 hours after transfection, the cell culture media (DMEM) was exchanged. 4 days after transfection, lentiviral particles were harvested by removing the supernatant from the well and aliquoting in 1.5 ml tubes.

After harvesting lentiviral particles, a 6 well plate was seeded with HEK293Ta cells at a seeding density of 3×10^5^ cells in DMEM supplemented with 8mg/ml Polybrene (Sigma). Each well was then transduced with either 5μl, 10μl, 50μl, 100μl, 200μl or no virus 6 hours after seeding when cells had annealed to plate. 24 hours after transduction, the cell culture media was then exchanged for DMEM supplemented with 3μg/ml Puromycin (Thermo). Selection was carried out for 3 days to select for successfully transduced cells.

### Fluorescence microscopy

All transfections were carried out with 1μl of PEI reagent in a total volume of 50μl of Opti-MEM. Base-editing reporter HEK293Ta cells were plated 2 days prior to transfection on a 24-well tissue culture plate with a seeding density of 5×10^4^ cells per well. Two days later, wells were subsequently transfected with 250ng of Target-AID (Addgene 131125) and 250ng of SmartGuide vector with or without 250ng of Opener vector or cells were transfected with 250ng of Control vectors with 250ng of Target-AID.

4 days after transfection, cells were stained with 5μg/ml Hoechst and were imaged using a Keyence BZ-X800 high-resolution imaging system.

### In-vitro transcription of SgRNA

All experiments were performed at 34 °C in a CFX96 Touch™ (BioRAD) thermal cycler, and activity monitored upon fluorescence level shifts. Transcription was initiated upon addition of 4 nM of dsDNA template (gblocks Gene Fragment, IDT) to a solution of 5 U.μL^-1^ of T7 RNA polymerase (NEB, M0251), 1 U.μL^-1^ of murine RNase Inhibitor (NEB M0314), 500 μM of rNTPs (NEB, N0450) and 10 nM (1x) of SYBR™ Green II RNA Gel Stain (Invitrogen) in the reaction buffer containing 40 mM Tris-HCl, 6 mM MgCl_2_, 1 mM 1,4-Dithiothreitol, 2 mM spermidine with a final volume of 20 μL and incubated for 6 to 8 hours. We work in low yield conditions to reduce the proportion of transcription byproducts (ref). After transcription, RNA was isolated by precipitation in isopropanol-ethanol 75% and resuspended in 10 μL nuclease free water (NEB, B1500) before quantification by BioDrop μLITE (Biodrop). Transcripts were then stored at - 25 °C. Cognate miRNAs were chemically synthetized from IDT (standard desalting), resuspended in 10 mM Tris-HCl (pH 8), 1 mM EDTA and stored at -25 °C prior to use.

### Fluorescence cleavage assay

200 nM of isolated smgRNA were injected in a pre-incubating solution of 200 nM Cas9 (S. *pyogenes*, NEB, M0386), 120 nM fluorescent beacon and 1 U.μL^-1^ murine RNase Inhibitor (NEB M0314) in the reaction buffer containing 50 mM Tris-HCl (pH 7.5), 5.5 mM MgSO_4_, 75 mM NaCl, 100 μg.μL^-1^ BSA (NEB, B9000S) to a final volume of 10 μL per condition.

### One-pot assay

Reunion of 4-6 nM of dsDNA template (gblocks^®^ Gene Fragment, IDT) to a solution of 5 U/μL of T7 RNA polymerase (NEB, M0251), 200 nM Cas9 (S. *pyogenes*, NEB, M0368), 120 nM fluorescent beacon, 1 U.μL^-1^ of murine RNase Inhibitor (NEB M0314), 500 μM of rNTPs (NEB, N0450) and 10 nM (1x) of SYBR™ Green II RNA Gel Stain (Invitrogen) in the reaction buffer -containing 50 mM Tris-HCl (pH 7.5), 5.5 mM MgSO_4_, 75 mM NaCl, 100 μg.μL^-1^ BSA (NEB, B9000S) to a final volume of 10 μL per smgRNA.

