## supplementary material for "License to cut: Smart RNA guides for conditional control of CRISPR-Cas9"

Supplementary Figure 1 detailed smartguide sequence and secondary structure

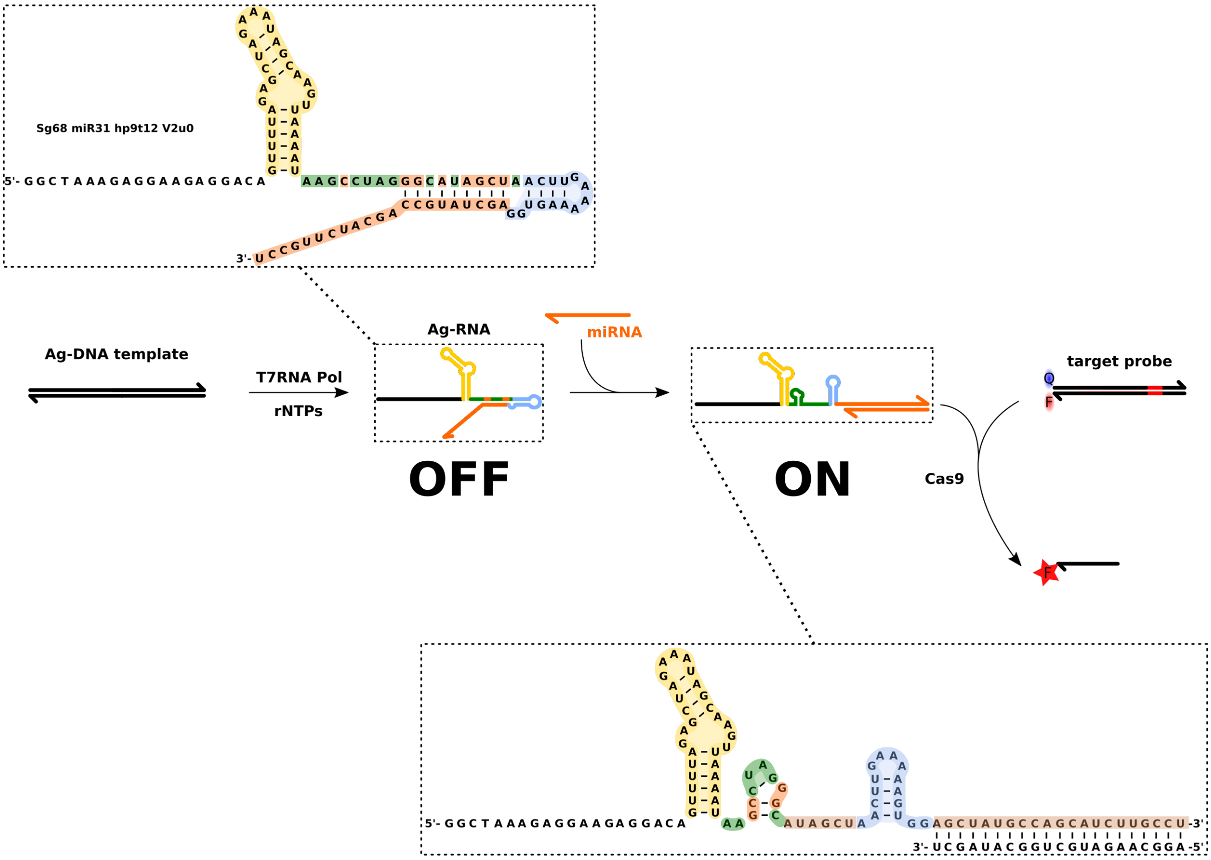

Supplementary Table X Primers used for construction of in-vivo Smartguide plasmid backbones

| **Primer Name** | **Sequence** |
|---|---|
| AA-sg1-BC-CS-eGFP-FW | TCCGGTGAATTCCCGTGTCCTCTTCCTCTTTAGTGGTGAGCAAGGGCGAGGAG |
| AA-eGFP-BamHI-RV | AGAATTGGATCCTTACTTGTACAGC |
| AA-MluI-U6-gib-FW | GAAACGGTACCACGGGCCAGATATACGCGTCGAGGGCCTATTTCCCATGA |
| AA-U6-sg1-scaf-short-RV | GCTATTTCTAGCTCTAAAACTGTCCTCTTCCTCTTTAGTGCGGTGTTTCGTCCTTTCCAC |
| AA-sg1-scaf-short-filler-FW | AGGACAGTTTTAGAGCTAGAAATAGCAAGTTAAAATAACGAGACGATGGCCTCCTCCGAGAAC |
| AA-filler-SpeI-gib-RV | GTGACCTCGAGCGGCCGCTCTAGAACTAGTGGAAAAAACGAGACGAAGTTCATCA |
| AvrII-1x-AUG-XFP-FW | TGAGGAGGCTTTTTTGGAGGCCTAGGGCCACCATGGTGAGCAAGGGCGAGG |
| BstB1-mCherry-RV | GGGCGTCGCTTGGTCGGTCATTTCGAATTACTTGTACAGCTCGTC |

Supplementary Table X Primers used for construction of SmartGuide plasmids or control plasmids by golden gate assembly

| **Plasmid** | **Forward Primer (5'-3')** | **Reverse Primer (5'-3')** |
|---|---|---|
| PB4-sg-31 | ATAAGCCTAGGGCATAGCTAACTTGAAAAAGTGGAGCTATGCCAGCATCTTGCCT | AAAAAGGCAAGATGCTGGCATAGCTCCACTTTTTCAAGTTAGCTATGCCCTAGGC |
| PB4-sg-31-3'hp | ATAAGCCTAGGGCATAGCTAACTTGAAAAAGTGGCACCGAGTCGGTGCAGCTATGCCAGCATCTTGCCT | AAAAAGGCAAGATGCTGGCATAGCTGCACCGACTCGGTGCCACTTTTTCAAGTTAGCTATGCCCTAGGC |
| PB4-sg-31-control | ATAAGCCTAGGGCATAGCTAACTTGAAAAAGTGGCACCGAGTCGGTGC | AAAAGCACCGACTCGGTGCCACTTTTTCAAGTTAGCTATGCCCTAGGC |
| PB4-sg-31-control (no nexus mutation) | ATAAGGCTAGTCCGTAGGCAACTTGAAAAAGTGGCACCGAGTCGGTGC | AAAAGCACCGACTCGGTGCCACTTTTTCAAGTTGCCTACGGACTAGCC |
| PB4-sg-31 (no nexus mutation) | ATAAGGCTAGTCCGTAGGCAACTTGAAAAAGTGGAGCTATGCCAGCATCTTGCCTACGGA | AAAATCCGTAGGCAAGATGCTGGCATAGCTCCACTTTTTCAAGTTGCCTACGGACTAGCC |
| PB4-sg-31-3'hp (no nexus mutation) | ATAAGGCTAGTCCGTAGGCAACTTGAAAAAGTGGCACCGAGTCGGTGCAGCTATGCCAGCATCTTGCCTACGGA | AAAATCCGTAGGCAAGATGCTGGCATAGCTGCACCGACTCGGTGCCACTTTTTCAAGTTGCCTACGGACTAGCC |
| PB4-sg-451 (no nexus mutation, 6bp toehold) | ATAAGGCTAGTCCGTATTCAACTTGAAAAAGTGGAACTCAGTAATGGTAACGGTTTACGGAA | AAAATTCCGTAAACCGTTACCATTACTGAGTTCCACTTTTTCAAGTTGAATACGGACTAGCC |
| PB4-sg-451 (no nexus mutation, 7bp toehold) | ATAAGGCTAGTCCGTAATCAACTTGAAAAAGTGGAACTCAGTAATGGTAACGGTTTACGGAA | AAAATTCCGTAAACCGTTACCATTACTGAGTTCCACTTTTTCAAGTTGATTACGGACTAGCC |
| PB4-sg-451 (no nexus mutation, 9bp toehold) | ATAAGGCTAGTCCGTAAACAACTTGAAAAAGTGGAACTCAGTAATGGTAACGGTTTACGGAA | AAAATTCCGTAAACCGTTACCATTACTGAGTTCCACTTTTTCAAGTTGTTTACGGACTAGCC |
| PB4-sg-synthetic (no nexus mutation, 8bp toehold) | ATAAGGCTAGTCCGTAGGCAACTTGAAAAAGTGGCGTTGATAGATGATGATGAATGGTAACTGGAGCCTACGGA | AAAATCCGTAGGCTCCAGTTACCATTCATCATCATCTATCAACGCCACTTTTTCAAGTTGCCTACGGACTAGCC |
| PB4-sg-451 (no nexus mutation, 9bp toehold) | ATAAGGCTAGTCCGTAGGCAACTTGAAAAAGTGGCGTTGATAGATGATGATGAATGGTAACTGGAGCCTACGGAA | AAAATTCCGTAGGCTCCAGTTACCATTCATCATCATCTATCAACGCCACTTTTTCAAGTTGCCTACGGACTAGCC |

| | | |
|---|---|---|
| PB4-sg-451 (no nexus mutation, 10bp toehold) | ATAAGGCTAGTCCGTAGGCAACTTGAAAAAGTGGCGTTGATAGATGATGATGAATGGTAACTGGAGCCTACGGACA | AAAATGTCCGTAGGCTCCAGTTACCATTCATCATCATCTATCAACGCCACTTTTTCAAGTTGCCTACGGACTAGCC |

Supplementary Table X Primers used for construction of SmartGuide opener plasmids

| **Plasmid** | **Forward Primer (5’-3’)** | **Reverse Primer (5’-3’)** |
|---|---|---|
| pcDNA3.1-miR-31-opener-mCherry | CAGATATACGCGTTTTCCCATGATTCCTTCATATTTGC | GGGCCCTCTAGAAAAAAAGCTATGCCAGCATCTTGCCTCGGTGTTTCGTCCTTTCCAC |
| pcDNA3.1-miR-451a-opener-mCherry | CAGATATACGCGTTTTCCCATGATTCCTTCATATTTGC | GGGCCCTCTAGAAAAAAAACTCAGTAATGGTAACGGTTTCGGTGTTTCGTCCTTTCCAC |
| pcDNA3.1-Synthetic-opener-mCherry | CAGATATACGCGTTTTCCCATGATTCCTTCATATTTGC | GGGCCCTCTAGAAAAAAACGTTGATAGATGATGATGAATGGTAACTGGAGCCTCGGTGTTTCGTCCTTTCCAC |

**Supplementary Table X** DNA template sequences for in-vitro experiments

| | guide | DNA template for in-vitro transcription |
|---|---|---|
| **Fluorescent reporters** | b2 bottom | ATTACGAATTCACCATGTCCTCTTCCTCTTTAGCCTAT**/Atto647N/** |
| | b2 top | **/BBQ-650/** ATAGGCTAAAGAGGAAGAGGACATGGTGAATTCGTAAT |
| | b3 bottom | ATTACGAATTCACCACTTTCCATTGGTCTCCTTCCTAT**/Atto 590/** |
| | b3 top | **/BMN-Q620/**ATAGGAAGGAGACCAATGGAAAGTGGTGAATTCGTAAT |
| **Matrix** | SmgRNA-98-3p | CACTACACTACTAACACACCACCAAA*TAATACGACTCACTATA*GGCTAAAGAGGAAGAGGACAGTTTTAGAGCTAGAAATAGCAAGTTAAAATAAGTCTAGTACTTTCCCAACTTGAAAAAGTGG**GGGAAAGTAGTAAGTTGTATAG** |
| | Positive control | CACTACACTACTAACACACCACCAAA*TAATACGACTCACTATA*GGCTAAAGAGGAAGAGGACAGTTTTAGAGCTAGAAATAGCAAGTTAAAATAAGTCTAGTACTTTCCCAACTTGAAAAAGTGG |
| | SmgRNA-451a | CACTACACTACTAACACACCACCAAA*TAATACGACTCACTATA*GGCTAAAGAGGAAGAGGACAGTTTTAGAGCTAGAAATAGCAAGTTAAAATAAGTCTAGTACTGAGTTAACTTGAAAAAGTGG**AACTCAGTAATGGTAACGGTTT** |
| | Positive control | CACTACACTACTAACACACCACCAAA*TAATACGACTCACTATA*GGCTAAAGAGGAAGAGGACAGTTTTAGAGCTAGAAATAGCAAGTTAAAATAAGTCTAGTACTGAGTTAACTTGAAAAAGTGG |
| | SmgRNA-3945 | CACTACACTACTAACACACCACCAAA*TAATACGACTCACTATA*GGCTAAAGAGGAAGAGGACAGTTTTAGAGCTAGAAATAGCAAGTTAAAATAAACCTAGGGTTGATATAACTTGAAAAAGTGG**ATATCAACCCTCTCCTATGCCCT** |
| | Positive control | CACTACACTACTAACACACCACCAAA*TAATACGACTCACTATA*GGCTAAAGAGGAAGAGGACAGTTTTAGAGCTAGAAATAGCAAGTTAAAATAAACCTAGGGTTGATATAACTTGAAAAAGTGG |
| | SmgRNA-21-5p | CACTACACTACTAACACACCACCAAA*TAATACGACTCACTATA*GGCTAAAGAGGAAGAGGACAGTTTTAGAGCTAGAAATAGCAAGTTAAAATAATCCTAGTGATGTTGAAACTTGAAAAAGTGG**TCAACATCAGTCTGATAAGCTA** |
| | positive control | CACTACACTACTAACACACCACCAAA*TAATACGACTCACTATA*GGCTAAAGAGGAAGAGGACAGTTTTAGAGCTAGAAATAGCAAGTTAAAATAATCCTAGTGATGTTGAAACTTGAAAAAGTGG |
| | SmgRNA-31 | CACTACACTACTAACACACCACCAAA*TAATACGACTCACTATA*GGCTAAAGAGGAAGAGGACAGTTTTAGAGCTAGAAATAGCAAGTTAAAATAAGCCTAGGGCATAGCTAACTTGAAAAAGTGG**AGCTATGCCAGCATCTTGCCT** |

| | | |
|---|---|---|
| | Positive control | CACTACACTACTAACACACCACCAAA*TAATACGACTCACTATA*GGCTAAAGAGGAAGAGGACAGTTTTAGAGCTAGAAATAGCAAGTTAAAATAAGCCTAGGGCATAGCTAACTTGAAAAAGTGG |
| | SmgRNA-122 | CACTACACTACTAACACACCACCAAA*TAATACGACTCACTATA*GGCTAAAGAGGAAGAGGACAGTTTTAGAGCTAGAAATAGCAAGTTAAAATAACACTAGATGGTGTTTGAACTTGAAAAAGTGGC**AAACACCATTGTCACACTCCA** |
| | Positive control | CACTACACTACTAACACACCACCAAA*TAATACGACTCACTATA*GGCTAAAGAGGAAGAGGACAGTTTTAGAGCTAGAAATAGCAAGTTAAAATAAGCCTAGGGCATAGCTAACTTGAAAAAGTGGTTTT |
| **Not gate** | SmgRNA-NOT-miR-98-3 | CACTACACTACTAACACACCACCAAA*TAATACGACTCACTATA*GGCTAAAGAGGAAGAGGACA**GGGAAAGTAGTAAGTTGTATAG**TACTAAGTTTTCCCAAGGCTAGTCCGTTATCAACTTGAAAAAGTGG |
| | Positive control | CACTACACTACTAACACACCACCAAA*TAATACGACTCACTATA*GGCTAAAGAGGAAGAGGACAGTTTTAGAGCTAGAAATAGCAAGTTAAAATAAGGCTAGTCCGTTATCAACTTGAAAAAGTGG |
| **notXandY gate** | NOT miR21 AND miR31 | CACTACACTACTAACACACCACCAAA*TAATACGACTCACTATA*GGCTAAAGAGGAAGAGGACAGTTTTAGAGCTA**TCAACATCAGTCTGATAAGCTA**TAGCAAGTTAAAATAAGCCTAGGGCATAGCTAACTTGAAAAAGTGG**AGCTATGCCAGCATCTTGCCT** |
| | Positive control (Sg68) | CACTACACTACTAACACACCACCAAA*TAATACGACTCACTATA*GGCTAAAGAGGAAGAGGACAGTTTTAGAGCTAGAAATAGCAAGTTAAAATAAGGCTAGTCCGTTATCAACTTGAAAAAGTGG |
| **XOR gate** | NOT miR21 AND miR31 | CACTACACTACTAACACACCACCAAA*TAATACGACTCACTATA*GGCTAAAGAGGAAGAGGACAGTTTTAGAGCTA**TCAACATCAGTCTGATAAGCTA**TAGCAAGTTAAAATAAGCCTAGGGCATAGCTAACTTGAAAAAGTGG**AGCTATGCCAGCATCTTGCCT** |
| | NOT miR31 AND miR 21 | CACTACACTACTAACACACCACCAAA*TAATACGACTCACTATA*GGCTAAAGAGGAAGAGGACAGTTTTAGAGCTA**AGCTATGCCAGCATCTTGCCT**TAGCAAGTTAAAATAATCCTAGTGATGTTGAAACTTGAAAAAGTGG**TCAACATCAGTCTGATAAGCTA** |
| | Positive control (Sg68) | CACTACACTACTAACACACCACCAAA*TAATACGACTCACTATA*GGCTAAAGAGGAAGAGGACAGTTTTAGAGCTAGAAATAGCAAGTTAAAATAAGGCTAGTCCGTTATCAACTTGAAAAAGTGG |
| **multiplex** | b2 miR21 | CACTACACTACTAACACACCACCAAA*TAATACGACTCACTATA*GGCTAAAGAGGAAGAGGACAGTTTTAGAGCTAGAAATAGCAAGTTAAAATAATCCTAGTGATGTTGAAACTTGAAAAAGTGG**TCAACATCAGTCTGATAAGCTA** |

| | | |
|---|---|---|
| | Positive control | CACTACACTACTAACACACCACCAAA*TAATACGACTCACTATA*GGCTAAAGAGGAAGAGGACAGTTTTAGAGCTAGAAATAGCAAGTTAAAATAA**TC**CTAGT**GATG**T**TGA**AACTTGAAAAAGTGG |
| | b3 miR31 | CACTACACTACTAACACACCACCAAA*TAATACGACTCACTATA*GGAAGGAGACCAATGGAAAGGTTTTAGAGCTAGAAATAGCAAGTTAAAATAAG**C**CTAG**GG**C**A**T**AGCT**AACTTGAAAAAGTGG**AGCTATGCCAGCATCTTGCCT** |
| | Positive control (Sg68) | CACTACACTACTAACACACCACCAAA*TAATACGACTCACTATA*GGAAGGAGACCAATGGAAAGGTTTTAGAGCTAGAAATAGCAAGTTAAAATAAG**C**CTAG**GG**C**A**T**AGCT**AACTTGAAAAAGTGG |

Supplementary Table XX Plasmid Maps

| **Name** | **Figure Panel** | **Benchling** |
|---|---|---|
| miR-31 smartguide with mutated nexus | C | https://benchling.com/s/seq-eI501UYMZjgI7ri5VIDL |
| miR-31 smartguide with mutated nexus+terminal stem loop | C | https://benchling.com/s/seq-N5AwTtYHdOBS0FEfpIBR |
| Mutated nexus+linker control | E | https://benchling.com/s/seq-uomQYt6NXUL2CJsVsC7W |
| WT nexus+mutated linker control | E | https://benchling.com/s/seq-H5LpOukSehTIiXjEEf6y |
| miR-31 smartguide with WT nexus | F | https://benchling.com/s/seq-Ght6X6bUZeCr9gzRZu0V |
| miR-31 smartguide with mutated WT+terminal stem loop | F | https://benchling.com/s/seq-xLwBwf8DB2b3AhOtyD9J |
| miR-451 smartguide, WT nexus, 6bp toehold | G | https://benchling.com/s/seq-sKVca4isJG4wbzwGybP8 |
| miR-451 smartguide, WT nexus, 7bp toehold | G | https://benchling.com/s/seq-rBr4FhIByiP1XcBJFgN3 |
| miR-451 smartguide, WT nexus, 9bp toehold | G | https://benchling.com/s/seq-WyMBKI7x2qQR0MI28 |

| | | |
|---|---|---|
| | | WZR |
| synthetic smartguide, WT nexus, 8bp toehold | G | https://benchling.com/s/seq-u9UgFYuKhuWNwHnsxfen |
| synthetic smartguide, WT nexus, 9bp toehold | G | https://benchling.com/s/seq-Y6iC6aNtWysIVfuyPwcF |
| synthetic smartguide, WT nexus, 10bp toehold | G | https://benchling.com/s/seq-bAjo3q3MaffvWUyDc3df |
